# Raf promotes dimerization of the Ras G-domain with increased allosteric connections

**DOI:** 10.1101/2020.07.15.205070

**Authors:** Morgan Packer, Jillian A. Parker, Jean K. Chung, Zhenlu Li, Young Kwang Lee, Trinity Cookis, Hugo Guterres, Steven Alvarez, MD Amin Hossain, Daniel P. Donnelly, Jeffrey N. Agar, Lee Makowski, Matthias Buck, Jay T. Groves, Carla Mattos

## Abstract

Ras dimerization is critical for Raf activation, yet Ras alone does not dimerize. Here we show that the Ras binding domain of Raf (Raf-RBD) induces robust Ras dimerization at low surface densities on supported lipid bilayers and, to a lesser extent, in solution as observed by size exclusion chromatography and confirmed by SAXS. Community network analysis based on molecular dynamics (MD) simulations show robust allosteric connections linking the two Raf-RBD D113 residues, located in the Galectin scaffold protein binding site of each Raf-RBD molecule and 85 Å apart on opposite ends of the dimer complex. Our results suggest that Raf-RBD binding and Ras dimerization are concerted events that lead to a high-affinity signaling complex at the membrane that we propose is an essential unit in the macromolecular assembly of higher order Ras/Raf/Galectin complexes important for signaling through the Ras/Raf/MEK/ERK pathway.

## Introduction

Ras GTPases are at the hub of signal transduction cascades that control cell processes such as proliferation, migration and survival, and their mutants appear in about a quarter of all human cancers, with poor treatment prognosis^1^. There are four main isoforms, each with specifically lipidated and highly divergent hypervariable regions (HVRs) that drive membrane localization and distinct biological outcomes^2^. The Ras G-domain is similar between the isoforms, KRas (4A and 4B), HRas and NRas, with sequence identical effector lobes (residues 1-86) and small differences in the allosteric lobe (residues 87-166)^3^. All four isoforms are purified as monomers, and HRas bound to the GTP analogue GppNHp has been shown by NMR to be strictly monomeric in solution^4^. Furthermore, full-length farnesylated and methylated KRas (KRas-FMe) does not form dimers by itself on supported membranes *in vitro*^5^. Paradoxically, indication that Ras functions through dimerization appeared in the early days of Ras research^6^ and Ras dimers have been detected at low levels in supported membranes^7-9^, in nanodiscs^10^, in cells^11^, in solution by NMR^12^ and mass spectrometry^13^. Importantly, dimerization has been shown to be essential for signaling through the Ras/Raf/MEK/ERK pathway in cells and in mice^14^. Computational studies of G-domain dimerization validated by some level of experimental evidence have identified three low-affinity dimerization interfaces in the absence of the Raf Ras-Binding Domain (Raf-RBD): an extended β-sheet formed at the effector lobe coinciding with the effector Raf binding interface^12,15^, and two overlapping interfaces involving α3-α4 or α4-α5 in the allosteric lobe^7,16,17^. Ras clustering on the membrane through these multiple interfaces may result in nanoclusters important for signaling, bringing together effector proteins and other components of the signaling machinery^18-20^. While nanoclusters may form through multiple Ras interfaces^18^, signaling through Ras/Raf occurs specifically through Ras dimers involving the α4-α5 interface^10,14^. Furthermore, the dimer of the Ras/Raf-RBD complex by itself is most likely not the full signaling unit, as scaffold proteins such as Galectins interact with Raf-RBD and are required for activation of the Ras/Raf/MEK/ERK pathway^21,22^.

The monomeric nature of the Ras G-domain in solution has led to skepticism about its role in dimerization, obscured by the fact that insertion of the C-terminal HVR region into the lipid membrane is important for dimerization in cells^11,23^. However, membrane insertion of the HVR is not sufficient for dimerization, as shown by experiments on supported membranes^5^. Here we use a combination of size exclusion chromatography (SEC) and small angle X-ray scattering (SAXS) experiments to show that the presence of Raf-RBD is sufficient to promote Ras G-domain dimerization in solution at levels detected by SEC in the absence of the HVR and membrane. Single molecule tracking experiments and fluorescence correlation spectroscopy demonstrate that robust dimerization of Ras on supported membranes requires Raf-RBD. Starting from a crystallographic model of the HRas/CRaf-RBD dimer (2 Ras/Raf-RBD interacting through the Ras α4-α5 interface), we use molecular dynamics (MD) simulations to show robust allosteric connections linking the Raf-RBD residues D113, located in the Raf-RBD binding site for Galectin^21^ and at the two opposing ends of the dimer of the Ras/Raf-RBD complex. Two independent 1 μs simulations starting from a model of the KRas/CRaf-RBD dimer with farnesylated KRas at the membrane show a stable dimer with allosteric linkages similar to those we obtain for the HRas/CRaf-RBD dimer. Our results, combined with those from the literature, suggest a model in which the dimer of the Ras/Raf-RBD complex is a key feature of a signaling platform that also includes the Galectin dimer.

## Results

### Raf promotes Ras G-domain dimerization in solution and on supported lipid membranes

While our Ras purification protocol yields a single peak in the final gel filtration step, addition of Raf-RBD to form the complex is accompanied by a higher molecular weight peak. Recent discussion of Ras dimers in the literature prompted us to investigate the contents of this higher molecular weight species that appears in the presence of Raf-RBD, as we found that the peak increases at high protein concentrations. The SEC profile for a protein solution containing the truncated G-domain of wild type KRas4B (referred to as KRas) bound to GppNHp in the presence of excess Raf-RBD shows three peaks that can be identified by SEC-SAXS data^24^ as the dimer of the Ras/Raf-RBD complex, the monomer of the Ras/Raf-RBD complex and excess Raf-RBD (Fig. 1). These results are in agreement with the molecular weights for the species of each of the three elution volumes determined by SEC based on a standard curve (Fig. 1 and Supplemental Materials Fig. S1). The SEC-SAXS results for HRas are very similar to those that we present here for KRas. In general, the relative intensities of the monomer and dimer peaks vary from one experiment to the next, depending on the Ras isoform, construct used (truncation at residue 166 or 173), protein concentration, the specific solution conditions, and experimental protocol. Raf-RBD is of paramount importance in these experiments, as Ras in the absence of Raf-RBD under similar experimental conditions does not form dimers (Supplemental Materials Fig. S1). Due to the presence of solvent exposed cysteine residues in the Ras/Raf-RBD complex (Raf-RBD C95 and C96 in particular), samples obtained from one of our SEC runs with the wild type constructs in presence of 1mM dithiothreitol (DTT) were submitted for mass spectrometry analysis. The results show that the observed Ras/Raf-RBD dimer peak does not contain disulfide bond crosslinks (Supplemental Materials Fig. S2). In further support of this, a SEC run with the Raf-RBD C95S-C96S double mutant, or with the wild type Raf-RBD in the presence of 100 mM DTT, shows that the dimer peak is still present (Supplemental Materials Fig. S3). The molecular envelopes generated from the SAXS data unequivocally support a dimeric Ras/Raf-RBD structure for the contents of the first major peak eluted from the SEC column (Fig. 1 and Supplemental Materials Fig S4), with excellent *χ*^2^ fits to either the crystallographic dimer (PDB ID 4G0N)^25^ or NMR dimer (PDB ID 6W4E)^10^ (with Raf-RBD added), both containing the α4-α5 interface (Supplemental Materials Fig. S5). This is consistent with the α4-α5 dimer interface having been established as the active dimer that promotes signaling through the Ras/Raf/MEK/ERK pathway^10,14^ and with signal inhibition by a monobody shown to bind at this interface^26^. The α4-α5 dimer interface is stabilized by hydrogen bonding and salt bridge interactions across the two Ras molecules (Fig. 2). Here we model the dimer based on our crystal structure of the HRas/CRaf-RBD complex generated by applying 2-fold crystallographic symmetry to the asymmetric unit (PDB ID 4G0N)^25^, supported by the appearance of the same α4-α5 interface in a large number of crystal structures^7,26^. In the recently published NMR data-driven model of KRas dimers on nanodiscs, the dimer affinity is low and the interface is highly flexible in the absence of Raf-RBD^10^. Although the average angle between helices across the dimer interface in the NMR model differs from that in the crystal structures, the key interacting residues are mostly the same.

**Fig. 1.**
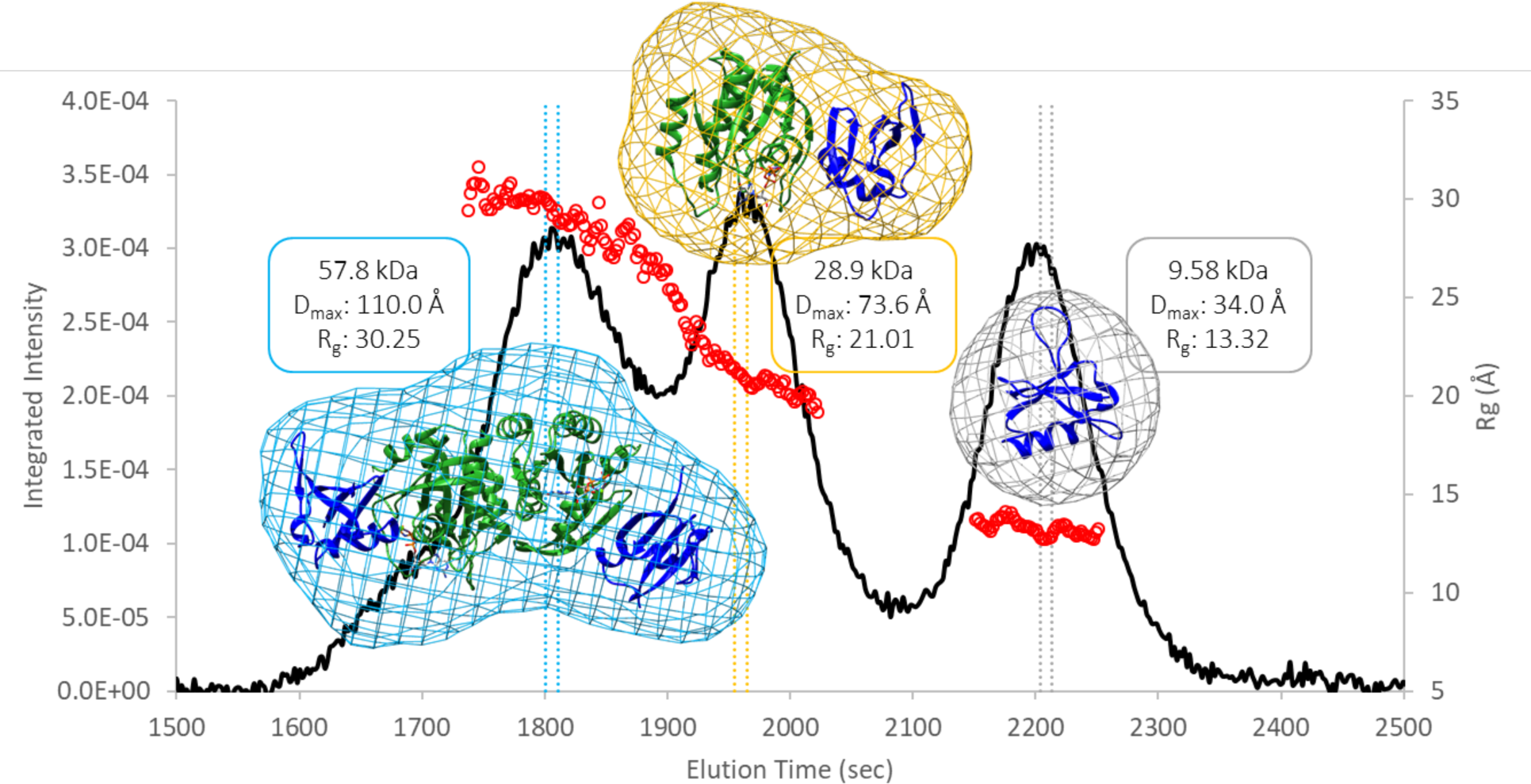
SEC-SAXS data collected for the three unique protein species that form in a solution containing the KRas-GppNHp G-domain and Raf-RBD. The black curve is the trace of the integrated X-ray scattering intensity (left axis), which correlates with protein concentration. The red data points correspond to the calculated radius of gyration (Rg, right axis) over elution time and indicate that the contents of each peak are monodisperse. The colored vertical bars through each peak indicate the data frames used to construct the corresponding molecular envelopes. Protein crystal structures were fit to the dimer (PDB ID 4G0N with dimer generated through a 2-fold crystallographic symmetry axis), monomer (PDB ID 4G0N asymmetric unity), and Raf-RBD (PDB ID 1RRB) envelopes using the volume fit function in Chimera.

**Fig. 2.**
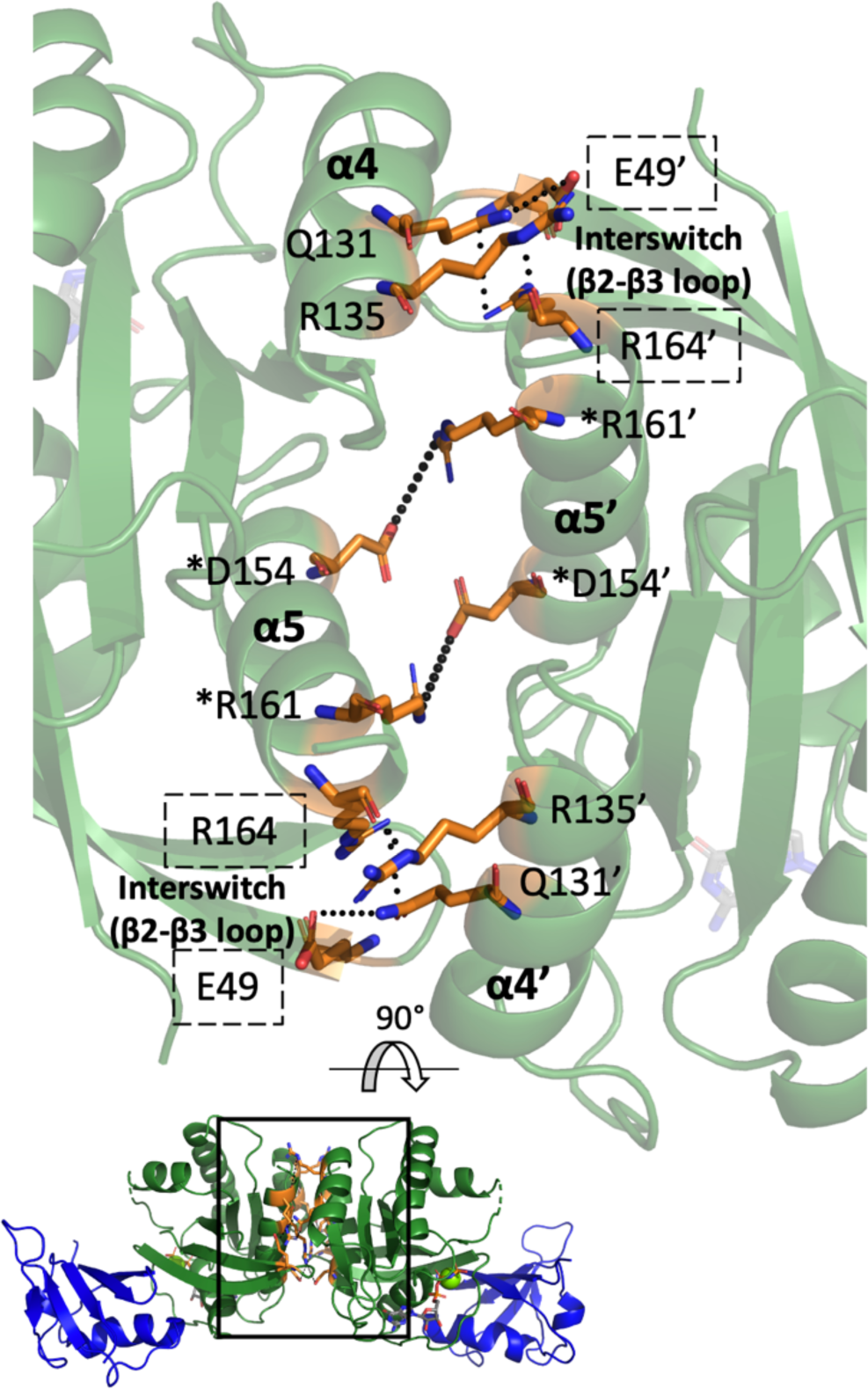
Interactions at the α4-α5 interface revealed by crystal structures and NMR models. (A) Interactions are shown for the interface found in our crystal structure of the Ras/Raf-RBD complex (PDB ID 4G0N with dimer generated through a 2-fold symmetry axis). Ras is in green, Raf-RBD is in blue. Residues at the dimer interface are in orange. The boxed interface area is rotated and magnified for clarity. Ras forms a symmetric dimer with corresponding residues in one of the monomers denoted by a prime (‘). Residues present within the crystallographic interface but not the interface observed by NMR have their names boxed by dashed lines. Residues denoted by a star (*;) have been validated *in vivo*. Only the key residue interactions are depicted for clarity.

On supported lipid bilayers (SLBs), the addition of Raf-RBD to GppNHp-bound Ras leads to a robust formation of complexes containing two Ras molecules (Fig. 3A,B). We detect this complex formation by the protein surface density-dependent decrease in diffusion, which can be used as an indicator for dimerization on a homogeneous membrane environment such as the one provided by SLBs^5,8,9^. Figure 3 shows full-length farnesylated and carboxy-methylated KRas (KRas-FMe)^27^ density-dependent changes in KRas diffusion with or without Raf-RBD measured by fluorescence correlation spectroscopy (FCS) and single molecule tracking (SMT). In the FCS measurements, the diffusion coefficient of KRas is unchanged across all observed surface densities, but is reduced from 4.5 to 2.3 µm^2^/s in the presence of Raf-RBD (Fig. 3C). A similar change can be seen in the SMT step size distributions (Fig. 3D). The change in the diffusion coefficient is identical to that due to dimerization by a crosslinker, suggesting that this complex contains two Ras molecules^5^. The data on SLBs (Fig. 3) indicate a 2-dimensional Kd in the order of tens of molecules per μm^2^, which corresponds to a very high affinity interaction likely due to allosteric effects leading to concerted binding of Raf-RBD and dimerization in the presence of the membrane. HRas shows a similar diffusion behavior (Supplemental Materials Fig. S6). The type and location of the fluorescent tag on Raf-RBD, necessary for the SLB experiments^28^, was found to be capable of disrupting the complex formation (Supplemental Materials Fig. S6B). While the addition of the SNAP tag (19.4 kDa) at the N-terminus removed the density-dependent decrease in diffusion, mCherry (28.8 kDa) on the C-terminus had no effect on complex formation. This is consistent with our model of the dimer, where the N-terminus of Raf-RBD is located close to the Ras/Raf-RBD interface and the C-terminus is in a position remote from the interfaces, where an added tag would not be expected to interfere with either Ras binding or dimerization. Overall, our experiments show robust dimerization of the Ras/Raf-RBD complex in supported membranes for both KRas and HRas.

**Fig. 3.**
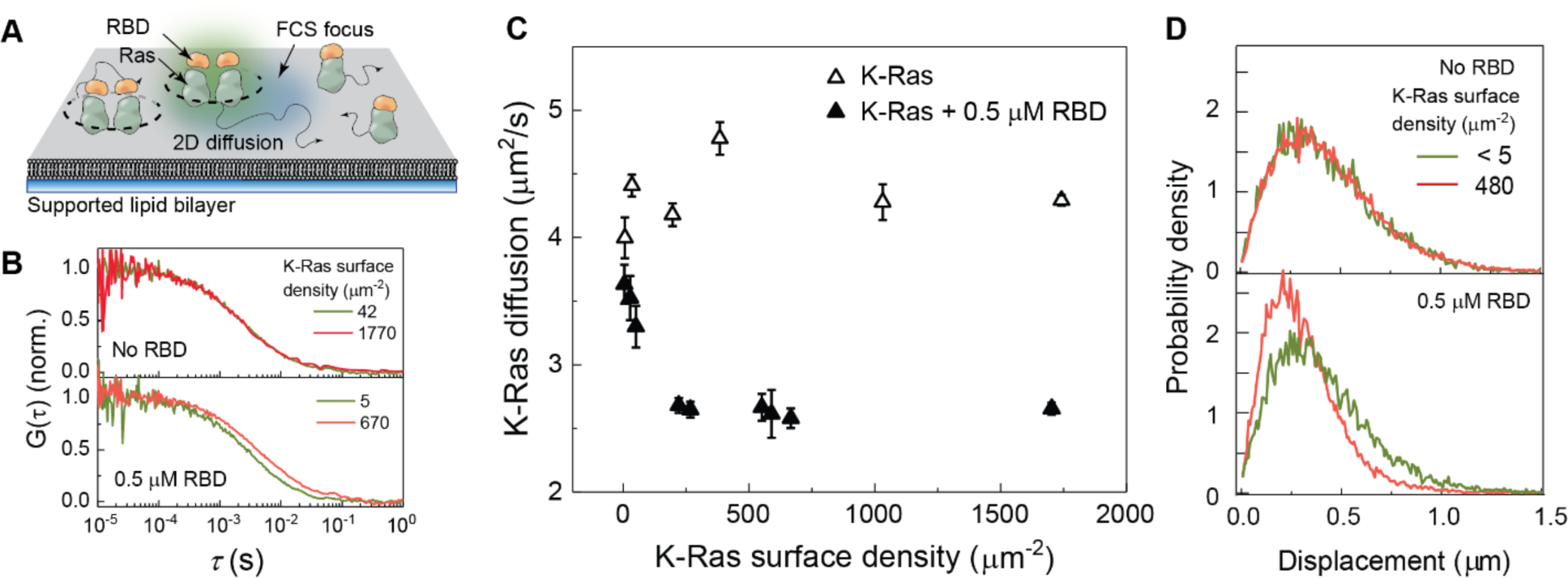
Ras diffusion measurements on supported lipid bilayers (SLBs). (A) The experimental setup on SLB. Farnesylated and methylated full-length KRas is allowed to spontaneously insert to the SLB, and Raf-RBD is introduced. In FCS, the surface density and the diffusion coefficient of fluorescent species diffusing together are measured. (B) FCS autocorrelation functions for KRas without (top) and with (bottom) 0.5 µM Raf-RBD for multiple KRas membrane surface densities. Only in the presence of RBD is there a Ras density-dependent change. (C) The apparent diffusion coefficients for KRas measured by FCS; the surface density-dependent decrease in diffusion indicates RBD-dependent oligomerization of Ras. (D) Single-molecule step size distributions without (top) and with (bottom) 0.5 µM Raf-RBD for KRas at low and high surface densities. SLBs were composed of 20% DOPS and 80% DOPC.

### Raf binding and dimerization of the Ras G-domain increase allosteric connections

Given our previous work showing that the dynamics of the Ras G-domain are affected by Raf-RBD through allosteric effects^25^, we performed 90 ns MD simulations on HRas, on the HRas/CRaf-RBD complex, and on the dimer of the HRas/CRaf-RBD complex to determine possible differences in allosteric communication between the Raf-RBD binding site and the dimer interface on Ras. HRas was used so that the simulations could be started directly from our crystal structures of HRas alone (PDB ID 3K8Y) and in complex with Raf-RBD (PDB ID 4G0N monomer and dimer). Dynamic network analysis^29^ of the protein trajectories using the Carma software package^30^ identifies correlated motion between protein residues. Visualizing these correlations reveals edges connecting the alpha carbon of residues, represented by spherical nodes, whose atoms are within 4.5 Å of one another throughout at least 75% of the simulation time, highlighting areas of correlated movements within a protein system^29^. These networks can be subdivided into communities of nodes that interact more with one another than with nodes of other communities, helping to identify allosteric networks. The communities calculated from the three simulations are shown in Figure 4. The simulation of Ras by itself shows six distinct communities, indicating six regions of the protein with relatively independent dynamics (Fig. 4A, left panel). Note that neither of the helices involved in Ras dimerization, helices 4 (yellow) and 5 (blue), are connected to each other or to the effector lobe of Ras, and the Ras active site is divided into three communities. In addition to determining communities of connected residues, it is possible to trace the edges connecting any two residues using optimal and suboptimal path calculations, where the optimal path shows the smallest number of nodes, or shortest path, between “source” and “sink” residues connected through long-distance interactions, and the suboptimal paths show all other connections differing by no more than 20 edges from the optimal path^29^. Taking our simulation of Ras-GTP by itself, we checked our previously identified allosteric network connecting the allosteric site residue R97 to the active site residue Q61 on switch II^31^ and found a total of 10 optimal and suboptimal paths connecting the two residues (Fig. 4A, right panel).

**Fig. 4.**
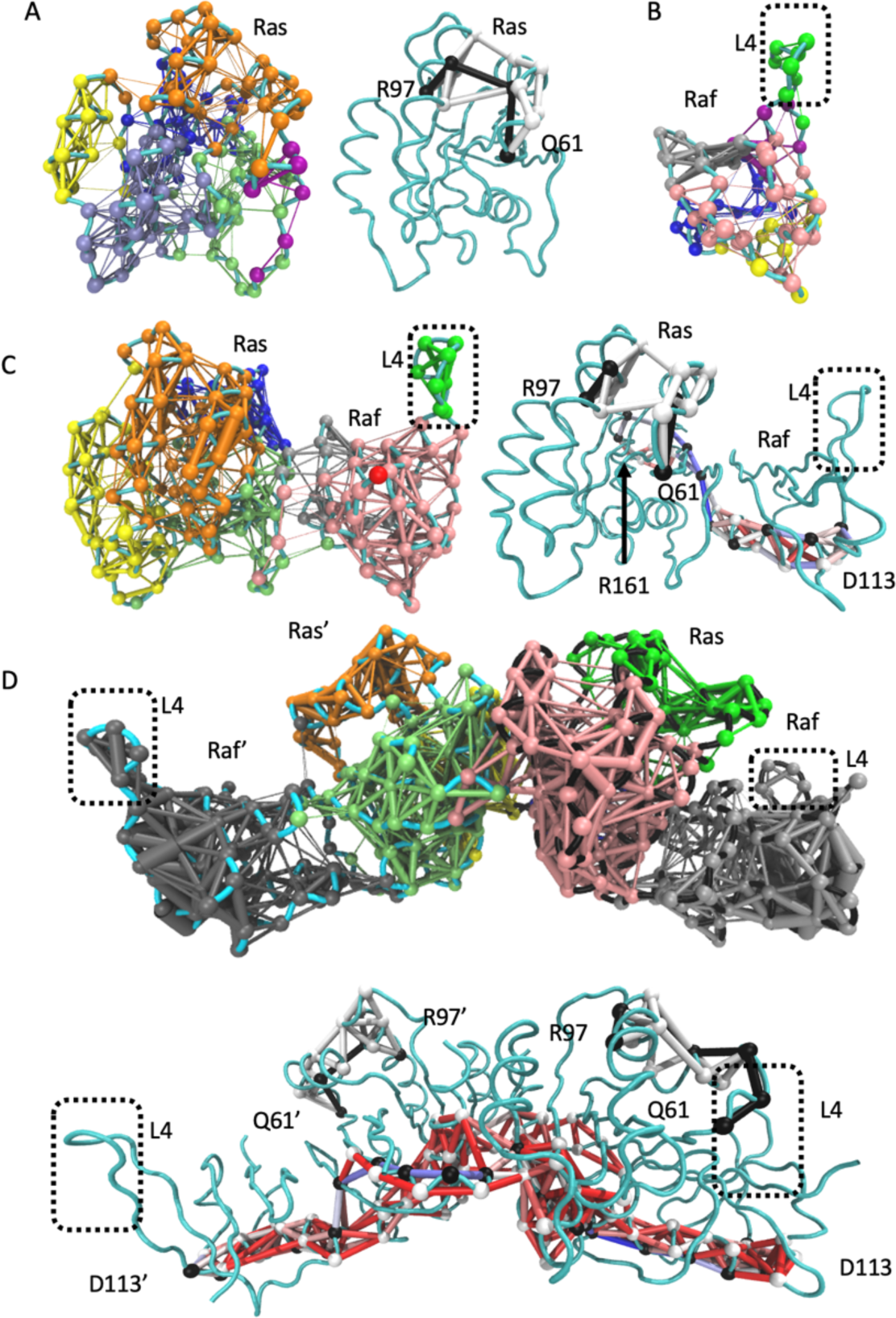
Community network and path analyses performed by Carma and visualized using NetworkView in VMD. (A) Ras alone, (B) Raf-RBD alone, (C) the monomer of the Ras/Raf complex, and (D) the dimer of the Ras/Raf complex. Left panels in A and C show community network analyses, with each community in a distinct color, and right panels show optimal (black nodes and edges) and suboptimal (white nodes and edges) path analyses between R97 and Q61. Paths between R161 and D113 in the Ras/Raf-RBD complex or D113 and D113’ in the dimer (black nodes, optimal; white nodes, suboptimal) are shown with all edges colored by the number of paths that cross them, from the highest colored red, to the lowest colored blue. In D the analogous panels are on top and bottom. Spheres are nodes centered on the alpha carbons of amino acid residues and sticks are edges calculated by Carma.

Raf-RBD by itself also has six communities and, as with Ras, this indicates various regions with relatively independent dynamics (Fig. 4B).

In the monomer of the Ras/Raf-RBD complex, Ras coalesces into four communities, with residues within the Ras active site establishing connections to helix 5 (green), Raf-RBD (pink), and the allosteric site (orange) (Fig. 4C, left panel). Helix 4 strengthens its connections across the allosteric lobe towards the Ras active site (yellow). Upon complex formation, Raf-RBD also experiences an increase in allosteric connections, with three major communities (Fig. 4C). For calculation of the optimal and suboptimal paths across the monomeric complex we chose R161 in helix 5 at one end and D113 on Raf-RBD at the opposite end of the complex. Ras R161 is in a critical position in helix 5 at the dimer interface (Fig. 2). Raf-RBD D113 is situated near loop 4 and has been previously identified for its long-distance contribution to the Ras and Raf-RBD interaction^32,33^. D113 is also in the binding site between Raf-RBD and the scaffold protein Galectin^21^. In the Ras/Raf-RBD monomer, commnication develops between Ras helix 5 across the complex interface to residues near Raf-RBD loop 4, which is in its own dynamic community, as it is in Raf-RBD alone (Fig. 4C)^25^. There are over 50 paths linking Ras R161 to Raf-RBD D113 in the monomer of the complex, while the number of paths linking allosteric site R97 and active site Q61 on Ras remains small.

Simulations of the dimer of the Ras/Raf-RBD complex show a robust increase in connectivity, with Ras having three communities and Raf-RBD linked in a single allosteric network from the interface with Ras all the way to loop 4, near D113 on the opposite side of the molecule (Fig. 4D, top panel). The increase in allosteric connections upon dimerization of the complex links the entire β-sheet core of Ras to both of the dimerization helices (pink and light green) and unifies helix 5 into a single community with the entire Ras active site (light green).

Despite the increase in allosteric connection from one end of the dimer to the other, the portion of Ras that undergoes a conformational shift upon ligand binding at the allosteric site remains in an isolated community in each protomer (orange and bright green). The two Ras/Raf-RBD complexes in the dimer are approximately symmetric, although some asymmetry is observed, perhaps due to stochastic motions over the course of the simulations. In the highly connected dimer complex, about 20 paths link the allosteric site and active site residues and over 300 paths exist between helix 5 residue 161 at the dimer interface and the opposite end of the complex at Raf residue D113 (Figure 4D, top panel). Residue D154, another key residue at the dimer interface, also has over 300 paths to the Raf residue D113. However, when the D113 residues in the two Raf-RBD molecules, at a distance of 85 Å apart on opposite ends of the Ras/Raf-RBD dimer, are used as “source” and “sink” nodes for allosteric connectivity, over 25,000 paths are identified, reflecting a strong allosteric linkage between the two ends of the dimer through the dimer interface, with R161 included in about 40% of the paths. The most traversed inter-Ras edge among the possible D113-D113’ paths involves residue E143 of one Ras molecule and residue D47 of the other, consistent with residue D47 of each Ras molecule belonging to the same community as helix 4 in the opposite Ras protomer (Fig. 4D, pink and green). Interestingly, E143 is part of the active site ExSAK motif, where it forms a salt bridge with loop 8 residue R123 immediately following the nucleotide binding NKxD motif^3^. Thus, the Ras/Ras interface is strongly connected to the Ras/Raf-RBD interface and the active site at switch I in the dimer of the complex, providing a venue through which Raf-RBD binding may modulate Ras helices 4 and 5 to form a high affinity dimer. From Ras loop 8, the allosteric paths go though Ras helix 1 near the active site, across the Ras/Raf-RBD interface to Raf-RBD helix 1, towards D113 near loop 4, spaning the entire base of the complex (Figure 4D, bottom panel). Overall, it is clear that allosteric connectivity and information transfer are significanly strengthened in the dimer of the Ras/Raf-RBD complex, as supported both by community network analysis and allosteric paths between connected nodes that span the entire complex (Fig. 4D). Interestingly, the connection between the active and allosteric sites remain relatively weak and outside of the strong allosteric network that links the two Raf-RBD ends in the dimer of the Ras/Raf-RBD complex.

### MD simulations of the KRas/CRaf-RBD dimer on the membrane

Because KRas is a major Ras isoform of interest in Ras mutant cancers, we modeled the KRas/CRaf-RBD dimer complex based on our HRas/CRaf-RBD dimer (PDB ID 4G0N) and performed two independent 1 μs MD simulations of the model dimer of the KRas/CRaf-RBD complex on an 80%:20% POPC:POPS membrane. The average structure of the complex from the simulations is shown in Fig. 5A. The part of the dimer containing the G-domains bound to Raf-RBD has an excellent fit to our KRas/Raf-RBD SAXS data obtained from the dimer SEC peak, with a *χ*^2^ of 1.04 (Supplemental Materials Fig. S7A). The fit for the average model from our 90 ns simulations of the HRas/Raf-RBD dimer to the KRas/Raf-RBD SAXS data has a *χ*^2^ of 1.21 (Supplemental Materials Fig. S7B). Overall, the dimer of the KRas/CRaf-RBD complex is well maintained on the model membrane, consistent with the experimental finding on SLBs (Fig. 3). The root mean square deviation (RMSD) of the KRas dimer from the starting structure shows variations less than 6 Å (Supplemental Materials Fig. 8A). The fluctuations of the KRas/CRaf-RBD dimer relative to the model membrane are limited as reflected by its orientation angle (Supplemental Materials Fig. 8B), with the Ras helices 3, 4 and 5 remaining roughly perpendicular to the membrane, as was previously observed for the NRas dimer simulation on the membrane in the absence of Raf^7^. The dimer of the KRas/CRaf-RBD complex lifts away from the membrane in comparison to previous simulations of a model of the KRas/CRaf-RBD-CRD monomer^34^. While there are extensive membrane contacts for the HVR, the KRas G-domain as well as Raf-RBD are mostly away from the membrane throughout the simulations (Supplemental Materials Fig. 8C, D), as major membrane interaction regions in KRas helices 4 and 5 observed for monomeric KRas are now occupied by the dimerization interface. Membrane interactions are occasionally seen for R73 in switch II and R102 and D105 in loop 7 at the allosteric site (Fig. 5B). These residues are predicted to be exposed in the dimer to membrane phospholipid headgroups for possible allosteric modulation of GTP hydrolysis in the presence of Raf^25,31^. There is a significant increase in fluctuation of the angle between the α4-α5 helices across the Ras dimer interface in the simulations of the KRas/Raf-RBD dimer in the presence of the membrane compared to what we observe for the HRas/Raf-RBD dimer in the absence of the membrane, although the average angle differs by less than 10° (Supplemental Materials Fig. S8E). The most notable change is weakening of interactions at the bottom half of the dimer interface on the KRas dimer complex, as Raf-RBD is drawn toward the membrane to form transient electrostatic interactions involving residues 101-109 in loop 4 (Fig. 5C). *In vivo*, the interactions between Raf-RBD and the membrane in the context of the dimer could be shielded with the binding of the scaffold protein Galectin at the site including loop 4 and extending to D113^21^, perhaps resulting in a more stable Ras dimer interface on the membrane.

**Fig. 5.**
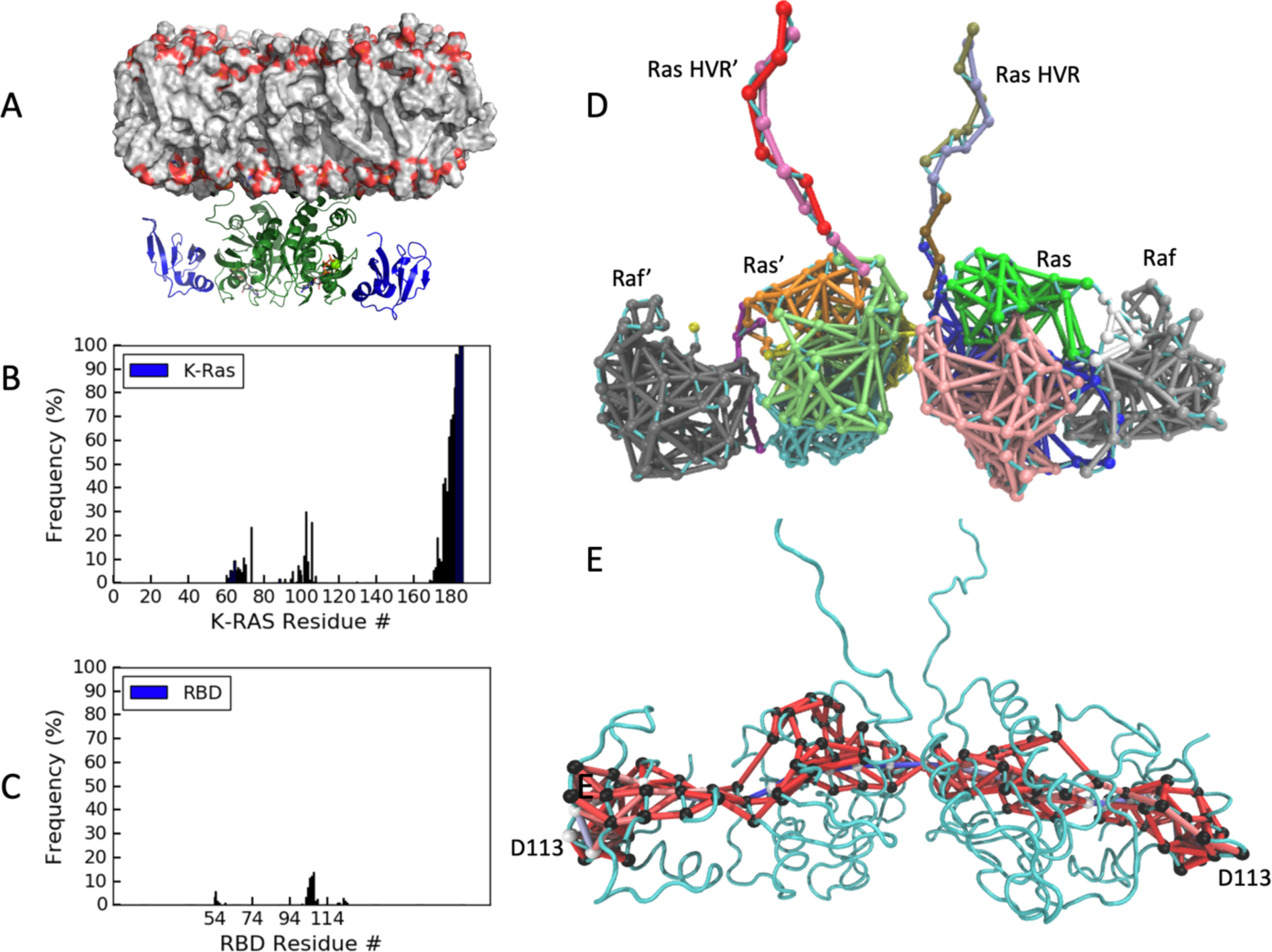
Dimer of the KRas/CRaf-RBD complex simulations at the membrane. (A) Average structure of the complex at a model membrane. The proteins are shown as ribbon diagram, with Ras in green and Raf-RBD in blue. The phospholipids on the membrane are colored by atoms with carbon in gray, phosphorous in orange and oxygen in red. (B-C) Frequency of KRas and Raf-RBD residues in the dimer coming within 4 Å of the membrane surface. (D) Community network analysis for the KRas/Raf-RBD dimer with communities colored as in Fig. 4D, top panel. (E) Optimal and suboptimal paths connecting Raf-RBD D113 residues on opposite ends of the complex. Nodes are white in the optimal path and black in the suboptimal paths, with all edges colored by the number of paths that cross through them as described in Fig.4.

In spite of the differences noted above, community network analysis and sub-optimal path calculations on simulations of the KRas/Raf-RBD dimer complex show similar patterns of allosteric connectivity as described for the HRas/Raf-RBD dimer (Fig. 5D). Once again incorporation of the Ras active site into the dimer interface communities was observed, with Raf coalescing into a single community of synchronous residues. There are minor communities observed in the switches of the KRas dimer complex not present in the HRas simulations, consistent with empirical data showing greater dynamics for KRas than for HRas in this region^35^. Importantly, the community containing the allosteric site and switch II (Fig. 5D orange and bright green) is segregated from the communities linking the two ends of Raf-RBD, as observed in the HRas/Raf-RBD simulations. Strong connections across the length of the Ras dimer linking residues D113 on the opposing Raf molecules are still present, although the optimal and suboptimal paths that cross the dimer interface traverse further up in the complex on KRas (Fig. 5E), closer to the membrane and further from loop 8, than observed for HRas (Fig. 4D). In the KRas dimer simulations, the edge between residues I139 and isoform-specific residue K165 at the top of helices 4 and 5 respectively, provide the most prominent connection across the dimer interface. Although interactions between residues I139 and Q165 are present in the dimer of the HRas/Raf-RBD complex, they are not part of a major allosteric pathway. Conversely the inter-switch loop 3 residues D47 and E49, which in the HRas/Raf-RBD dimer are at the center of allosteric pathways across the interface, interact with isoform specific helix 4 residue K128 (R128 in HRas) and with R135, but are not part of the calculated optimal and suboptimal paths in the KRas/Raf-RBD dimer. These differences between the KRas and HRas simulations could stem from isoform-specific residues at the dimer interface (K/R128, D/E153, K/Q165 KRas/HRas residues) or elsewhere, from the presence of the longer helix 5 leading to the HVR in the KRas simulations, or from the fact that the membrane is present for the KRas simulations but not for the HRas simulations. Furthermore, these details may change in the presence of Galectin. Importantly, regardless of whether the simulations include KRas or HRas, the community network analyses described above (Fig. 4 and Fig.5) show strong allosteric connections that link the Galectin-binding D113 residue near loop 4 in one Raf-RBD molecule to D113 on the other across the two Ras molecules.

## Discussion

While signaling through Raf is one of the major pathways by which Ras promotes cell proliferation and its mutants result in oncogenic phenotypes, the molecular mechanisms associated with Ras activation of Raf remain obscure. Here we advance the mechanistic understanding of Raf activation by Ras with evidence that Raf and the membrane act in concert to promote dimerization of the Ras/Raf complex, and show that dimerization dramatically increases allosteric connections linking the two Raf-RBD molecules at opposite ends of the dimer formed by the α4-α5 interface across the two Ras protomers. Having previously shown that HRas and KRas have distinct conformational preferences^3,35^, here we compare dynamic network analysis from 90 ns simulations of the HRas/CRaf-RBD dimer started directly from the crystal structure of the complex with longer simulations on the membrane, starting from a model of the KRas/CRaf-RBD dimer generated from the HRas/CRaf-RBD structure. In spite of the different setups, we found similar patterns of allosteric connections in our simulations (Fig. 4 and Fig. 5). Furthermore, SEC-SAXS experiments yielded similar molecular envelopes for dimers containing KRas/Raf-RBD and HRas/Raf-RBD (data not shown), the SLB experiments gave similar results for both isoforms (Fig. 3 and Supplemental Materials Fig. S6), and the dimerization interface for the KRas/Raf-RBD dimer modeled on HRas/Raf-RBD is stable throughout the 1 μs simulations, yielding a structure with excellent agreement to both KRas/Raf-RBD (*χ*^2^ of 1.04, Fig S7A) and HRas/Raf-RBD (*χ*^2^ of 1.26, data not shown) SEC-SAXS data sets. Together, these data point to a similar mechanism between the two Ras isoforms for signaling through the Ras/Raf/MEK/ERK pathway, which we expect would also extend to the NRas isoform.

Computational work has identified weak dimer interfaces involving the α3-α4 or α4-α5 Ras allosteric lobe helices^16^. Which interface is involved in signaling through Raf has until recently remained an open question, primarily because mutations in both helices 3^36^ and 5^14^ result in attenuated signaling through Ras/Raf. However, we have previously shown that a single mutation can have global allosteric effects on the structure and dynamics of the HRas/CRaf-RBD complex^25^. Additionally, given the long-range allosteric connections that we have demonstrated here through our community network analysis based on MD simulations, it is likely that mutations can affect dimerization without being at the interface. In general, given the highly allosteric impact of the Ras/Raf-RBD complex formation on both Ras and Raf-RBD^25,37^, great caution is needed in correlating the effects of specific mutations, or chemical shift perturbations obtained by NMR between bound and unbound species, to the locations of binding sites on Ras. The recent NMR structure of the KRas dimer on a nanodisc (PDB ID 6W4E) unequivocally shows that the dimer forms through the α4-α5 helical interface^10^, consistent with its prominent appearance in crystal structures of Ras^7,26^ and the stability of this interface in our simulations.

This interaction is weak in nanodiscs, with a Kd of 530 μM in the absence of Raf-RBD^10^, which is most likely observed due to the small membrane area on the nanodisc, leading to a crowded environment for the two Ras molecules. An analogous crowding situation occurs in Ras crystals, explaining the appearance of the dimer as previously tabulated^26^. In contrast, no dimerization is observed for Ras in the absence of Raf-RBD in SLBs^5^, which provide a larger area through which Ras molecules can diffuse.

It has been previously observed that un-complexed Ras has several regions of correlated motion, such as those between the C-terminal end of helix 3 and the switch regions^38,39^, and that the binding of Raf-RBD affects the dynamics of the entire monomer of the complex^25^. The present analysis identifies linkages between the dimer interface and the Raf-RBD loop 4 region in complexes with both HRas and KRas, in addition to further validating the connection between R97 in the allosteric site and Q61 in switch II, which we have previously studied^25,31^. The community on Ras that includes the calcium-binding allosteric site involving loop 7, which in our simulations transiently interacts with the membrane (Fig. 5B), remains relatively unchanged with binding of Raf-RBD and dimerization of the complex. However, while the allosteric connections linking the dimer interface in Ras to loop 4 in Raf are moderate in the monomer of the complex, a remarkable enhancement of allosteric connection occurs in the dimer of the Ras/Raf-RBD complex, linking residue D113 near loop 4 in both Raf-RBD molecules at the extreme ends of the complex. In this context, helices 4 and 5 at the dimer interface are intimately connected to the active site through the NKxD and ExSAK nucleotide-binding motifs, with helix 4 forming additional connections with loop 8 and the N-terminal end of helix 3, and helix 5 connecting to the inter-switch region and the β-sheet core. Overall, the major communities in Ras separate elements associated with GTP hydrolysis that we propose is promoted by calcium in the context of the dimer^25,31^, from those across the Ras dimer and Ras/Raf interfaces important for activation of Raf kinase. It is not surprising that these two separate functions, signaling and GTP hydrolysis, would be decoupled in this system.

Two recent cryo-EM structures of full-length Raf in complex with the scaffold protein 14-3-3^40^ and in complex with both 14-3-3 and MEK^41^ reveal details of activation at the Raf kinase/MEK level, but in the absence of Ras show disordered Ras binding domains, even as the kinase itself is in its active dimeric form. The fact that Raf kinase can dimerize in the absence of Ras points to a Ras function other than to simply promote kinase domain dimerization. When not in complex with Ras, the Raf cysteine rich domain (Raf-CRD) functions to maintain autoinhibition in the absence of active Ras, as revealed by the cryo-EM structure of the full-length inactive kinase, in which Raf-CRD is ordered and nestled between 14-3-3 and the Raf kinase domain^41^. This autoinhibition is released when bound to Ras, allowing for activation of the kinase, however, it is not clear that Ras dimerization is necessary for this process. Here we propose that a major function of Ras dimerization upon Raf binding is to couple with Galectin dimers to transform these protein complexes into an allosterically connected robust signaling platform (Fig. 6). The strong allosteric connection between the opposite extremes of the Ras/Raf dimer presents a unified complex that could be important for effective activation of signaling through Ras/Raf/MEK/ERK, particularly in the context of interactions with scaffold proteins such as Galectin, which is itself a homodimer critical for signaling through Ras/Raf^42,43^. Raf residue D113 is in the proposed binding site between CRaf-RBD and Galectin-1^21^, and is implicated in allosterically promoting Ras/Raf binding by at least three research groups using independent methods^21,44,45^. The importance of residue D113 to the signaling complex is further supported by the robust allosteric connections of this residue to critical Ras dimerization elements in helices 4 and 5. Coupling of the dimer of the Ras/Raf-RBD complex to the Galectin dimer through residue D113 in the loop 4 region could form a kinetic proofreading platform similar to that observed for the LAT/Grb2/SOS complex^28,46-48^.

**Figure 6.**
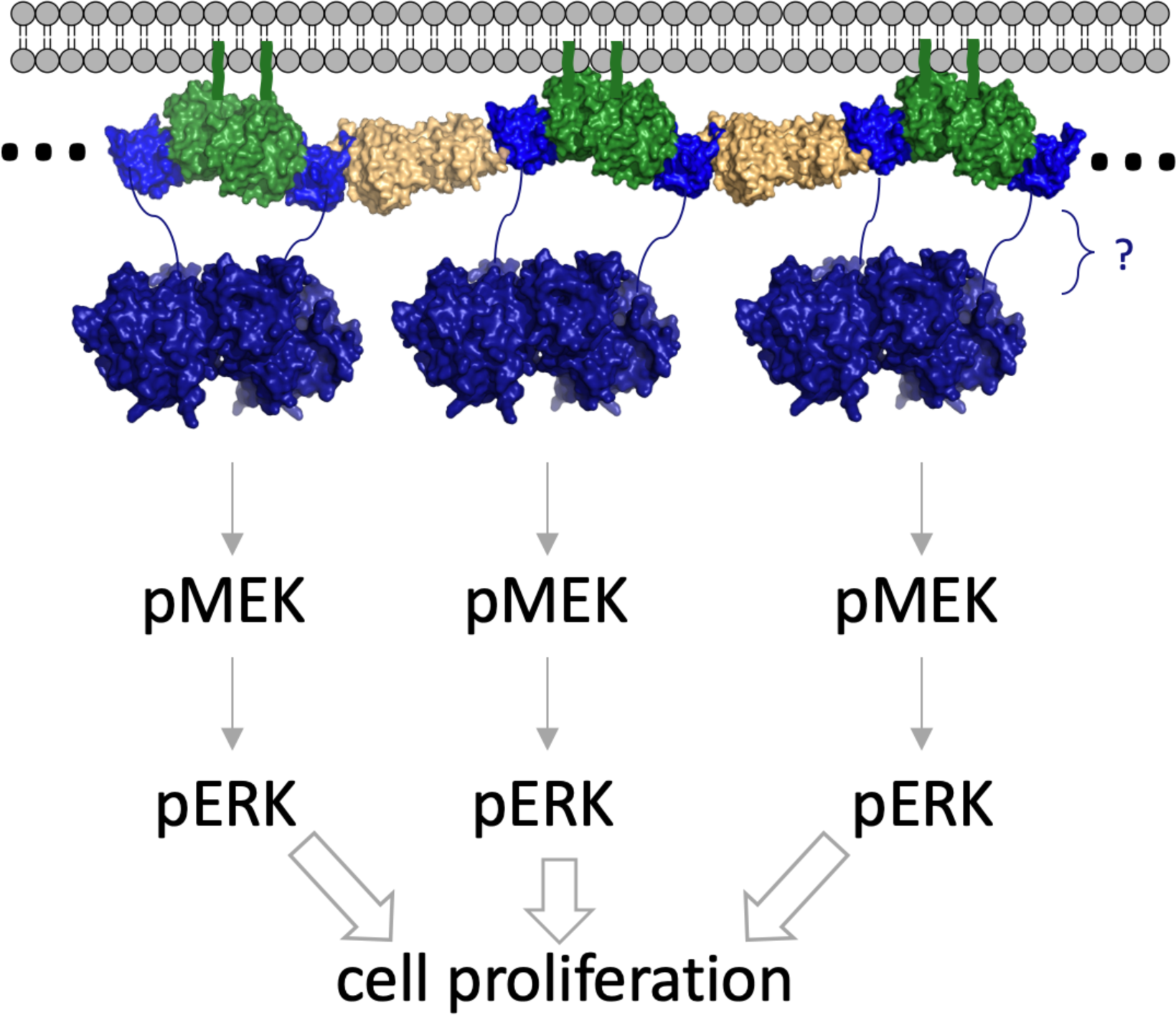
Multiprotein assembly proposed for signaling through the Ras/Raf/MEK/ERK pathway. Complexes involving Ras (green), Raf-RBD (blue), and Galectin dimers (beige) at the membrane would result in the generation of protein networks that result in synchronized activation for signal amplification and kinetic proofreading. The Raf-CRD and the serine/threonine rich region that follows, situated before the kinase domain (residues 132-340 in CRaf) are indicated by a blue line with a question mark to the right of the figure. The composite figure was made with structures for HRas/CRaf-RBD (PDB ID 4G0N), the Galectin-1 dimer (PDB ID 3W58) and the Raf-kinase domain dimer (PDB ID 3OMV).

Biocondensates, consisting of concentrated multiprotein assemblies forming distinct fluid structures that separate from surrounding areas, are ubiquitous across signal transduction networks^49^. Furthermore, single particle tracking (SPT) experiments^50,51^, most recently coupled with photoactivated localization microscopy (SPT-PALM) and detailed trajectory analysis^52^, have shown that activated Ras proteins in live cells are found on the membrane in mobile and immobile phases, the latter being consistent with the predicted formation of large macromolecular assemblies that cannot freely diffuse on the membrane. Recent work with the transmembrane receptor LAT and its cytosolic binding partners Grb2 and the Ras guanine nucleotide exchange factor SOS, have shown that these proteins form polymer-like assemblies on supported lipid bilayers^46^, with restricted mobility. By applying properties of polymer physics to this protein system, it becomes apparent that the proteins bound to several partners within the signaling competent complex form a gel-like phase at the membrane that crowding effects and Brownian motion fail to describe. Other interactions with the membrane from within these condensates, for example involving membrane receptors^53^, SOS^54-56^ or PLCγ^57^, further restrict molecular mobility and may influence Ras. Increasing connectivity with dimerization of the Ras/Raf complex could allosterically prime the Galectin binding site on Raf-RBD, to form a multivalent protein complex on the membrane corresponding to the immobile species observed in the SPT-PALM experiments^52^. This platform of synchronized activated signaling proteins (Fig. 6) could be essential for effective activation of the Ras/Raf/MEK/ERK signaling cascade, ensuring signal amplification and avoidance of random misfiring due to isolated encounters, in analogy to the LAT/Grb2/SOS platform. In this scenario, isolated Ras/Raf dimers would not be effective signaling agents, explaining the requirement for Galectin in the activation of the pathway^21,43^. Scaffold proteins are known to be hubs for the control of cell signaling^58^ and are often involved in biomolecular condensates associated with function either on or off membranes^49^. Our proposed model puts Ras dimerization at the center of such an organizational platform promoted by the binding of Raf-RBD and recruitment of Galectin for kinetic proofreading and signal amplification, leading to activation of the critical kinases in the mitogenic signal transduction pathway.

Ras by itself on the membrane forms nanoclusters consisting of oligomers with several Ras molecules interacting through multiple weak binding interfaces^16,23^. Given the robust high-affinity dimerization that we observe on SLBs in the presence of Raf-RBD, we propose that binding of Raf to Ras-GTP on the membrane allosterically modulates the α4-α5 interface such that it predominates in the signaling active dimer of the complex. This results in a dramatic increase in allosteric connectivity linking the Galectin-binding D113 residue on Raf-RBD near loop 4 at the two extremes of the dimer, which we propose becomes primed to interact with the Galectin dimer to form a robust and synchronized signaling platform. The present work provides a conceptual leap forward by revealing the relationship between Ras and Raf-RBD and the high allosteric connectivity between them, leading to the prospect that dimerization is a critical step in forming the signaling complex for synchronized activation of a large number of Raf kinase molecules.

## Methods

### Ras and Raf-RBD expression and purification

The G-domains of KRas or HRas proteins containing residues 1-166 or 1-173 were purified as previously described^59^. After nucleotide exchange to replace GDP with the GTP analogue GppNHp^60^, protein was loaded into a 5 mL sample loop followed by injection onto a HiTrap QHP 5 mL anion exchange column (GE Lifesciences). Ras was eluted with a linear 0–25% gradient of buffer B with composition described in the published protocol^59^. The peak fractions were analyzed by SDS-PAGE, and fractions containing Ras-GppNHp were pooled and concentrated to ∼20 mg/mL and flash-frozen in liquid nitrogen for storage at -80°C until needed. CRaf-RBD containing residues 52-131 was expressed and purified as previously described^3^.

### Ras-Raf-RBD Dimerization observed in solution by Size Exclusion Chromatography (SEC)

The appropriate Ras protein was combined with a 4-fold molar excess of Raf in stabilization buffer [20 mM HEPES pH 7.5, 20mM MgCl2, 50mM NaCl, 1mM DTT and 2% glycerol] and concentrated to ≤ 100 µL at 4°C. The solution was then spin-filtered through a 0.2 µm filter for injection onto a pre-equilibrated Superdex 200 Increase 10/300 GL column (GE Healthcare) for standard gel filtration analysis with a 0.5 mL/min flow rate. To reduce protein loss during injection onto the column, a 500 µL Hamilton syringe with large hub removable 22-gauge needle was used to inject the protein into a 100 µL sample loop via an INV-907 fill port. Peak integration was performed for the three major peaks, dimer K-Ras/Raf-RBD complex, monomer K-Ras/Raf-RBD, and excess Raf, using a zero baseline and the Unicorn peak integration function. Integration windows were adjusted to prevent inclusion of higher order oligomers that may be present in the sample mixture. The sum of the complex dimer and monomer peaks was taken as 100% complex and the respective percentage of dimer and monomers were calculated as the contribution of each peak to the total.

### Size exclusion chromatography in-line with small angle X-ray scattering (SEC-SAXS)

Samples for SEC-SAXS were prepared according to the dimerization assay protocol above, and flash frozen once concentrated for storage and transportation. Data were collected at the Cornell High Energy Synchrotron Source (CHESS) at Cornell University (Ithaca, NY). Standard size exclusion chromatography was run in stabilization buffer as above, where the eluted sample was subjected to constant light scattering data collection using a flow cell in-line with the X-ray beam ^61^. Because the elution profile matches exactly that of the standard Ras/Raf-RBD dimerization assay, the three peak identities for the SEC-SAXS chromatograms were known *a priori*. For each species (dimer Ras/Raf-RBD, monomer Ras/Raf-RBD, and Raf-RBD), five 2-second frames of data were combined (10s total exposure time) and buffer subtracted against five 2-second frames corresponding to the column equilibrated with stabilization buffer. The ATSAS program package was used for initial SAXS data analysis, including Guinier and Distance Distribution analysis, and envelope generation ^62,63^. SUPCOMB was used to align the molecular envelope generated using the SEC-SAXS data with the crystal structure PDB file (4G0N), with a very good NSD (Normalized Spatial Discrepancy) value of near 1^64^. Chimera was also used for alignment of the models in the SAXS-generated envelopes^65^. A direct comparison of the rigid PDB model with the SAXS data, yielding the reported *χ*^2^ values in Figures S5 and S7 was performed in FOXS^66^.

### Mass spectrometry

Samples eluted from the SEC peak corresponding to the dimer of the Ras/Raf-RBD complex in the presence of 1 mM DTT were prepared following a previously described protocol^67^. Briefly, protein from the appropriate pooled SEC fractions was precipitated using a volume ratio of 1:1:4:3 of protein:chlororform:methanol:water. The supernatant was removed and the precipitate washed with the addition and removal of another four parts methanol. Pellets were solubilized in cold (−20 °C) 80% formic acid and diluted to the original volume with HPLC-grade water. Intact protein LC-MS was performed using an H class Acquity Ultra High Pressure Liquid Chromatography (UPLC) system coupled with a Xevo G2-S Q-TOF mass spectrometer (Waters Corp, Milford, MA) as previously described^67^. Briefly, reversed phase chromatography was employed for separation (Acquity UPLC protein BEH C4 300 Å pore size, 1.7 µm particle size, 100 mm bed length, 2.1 mm ID x 100 mm) with 95% water/ 5% acetonitrile with 0.1% formic acid as solvent A and 95% acetonitrile/ 5 % water with 0.1% formic acid as solvent B.

### Fluorescence correlation spectroscopy (FCS)

FCS measurements were performed on a home-built confocal system integrated into an inverted microscope. The experimental methods have been published previously ^9^. The light source was a pulsed (100 ps at 20 MHz repetition rate) supercontinuum laser (NKT Photonics). The average excitation power for a typical FCS measurement was 0.5 µW for 488 ± 5 nm. The fluorescent signals were collected by the objective and passed a 50-µm pinhole detected by avalanche photodiode detectors (Hamamatsu), and processed by a hardware correlator (Correlator.com). To calibrate the spot size of the focus, a bilayer with a known surface density of fluorescent lipids, BODIPY-FL-DHPE for 488 nm, was measured, which consistently yielded the radii of 0.20 ± 0.01 µm. Each signal was focused into 0.15 × 0.15 µm avalanche photodiode elements (Hamamatsu), and subsequently processed by a hardware correlator (Correlator.com). The resulting autocorrelation *G*(*τ*) traces were fit to two-dimensional Gaussian diffusion model,

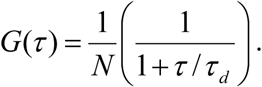

### Single-molecule tracking (SMT)

The experimental methods for smTIRF has been described previously^56^. TIRF images were acquired using a Nikon Eclipse Ti inverted microscope equipped with a 100× 1.49 NA oil immersion TIRF objective and an Andor iXon EMCCD camera. 488-nm and 637-nm diode lasers (Coherent Inc.) were used as illumination sources for TIRF imaging. The laser intensity was set to 0.5 and 15 mW at the objective for a 488 and 637 nm laser, respectively. In order to track KRas at the single molecule level in a wide range of surface density, 100 pM of Alexa 647-GppNHp-loaed KRas was mixed with various concentrations of eGFP-KRas loaded with nonfluorescent GppNHp (typically up to 50 nM). TIRF intensity of eGFP channel was used to estimate the overall density of KRas and Alexa 647-GppNHp-loaded KRas was imaged for single molecule tracking. The images were acquired at 20 ms exposure with no delay time. Particle localization and trajectory liking were done with TrackMate (ImageJ plugins)^68^. The step size distribution was calculated in Igor Pro (WaveMetrics).

### Molecular Dynamics Simulations

MD simulations of 90 ns production run were performed for Ras/Raf-RBD monomer and Ras/Raf-RBD dimer, where both structures were started from coordinates with PDB ID 4G0N^25^. The simulations were conducted at the Northeastern Discovery Cluster (http://www.northeastern.edu/rc). In each of the PDB files, the GTP analogue, GppNHp, was modified to form GTP by replacing the β-γ-bridging nitrogen atom with oxygen. Calcium acetate molecules were left at the allosteric site. All the crystallographic water molecules were included in the simulation. Each protein construct was additionally solvated by TIP3P waters with 150 mM NaCl (plus net charge neutralizing ions) for a total of 28,808 and 61,101 atoms for the monomer and dimer, respectively. Each system was minimized for 5,000 frames then equilibrated to 300 K. A 1 fs time step was used for the first 30 ns, then increased to 2 fs for the remaining 60 ns. Standard parameters were used with Particle-Mesh Ewald (PME) method for long range electrostatic interaction, a cut-off of 1.1 nm for the van der Waals (vdW) potential and short electrostatic interaction, SHAKE algorithm for constraint of all covalent bonds to hydrogen, and a Langevin thermo- and baro-stat at 300 K and 1 bar were used, respectively. The simulations were prepared using VMD and performed on NAMD software^69^. The CHARMM force field was used for the simulation^70^.

Molecular models of the 2:2 KRas/Raf-RBD dimer was constructed using available crystal structures (PDB 4DSO for KRas4B and 4G0N for the RBD; 4G0N is a complex of H-Ras/Raf-RBD which showed the dimer in the crystal lattice, onto which KRas4B was modeled. The RBD was then homology docked onto each side of the K-Ras4B dimer). The complex was anchored into a membrane of 280 POPC and 70 POPS (80%:20%). Two POPC lipid molecules were removed on the leaflet where KRas is anchored, in order to relax the membrane tension. The system was charged at pH 7.0 (using amino acid protonation states, and HSD, hydrogen on delta position for a neutral histidine) solvated by TIP3P water with 150 mM NaCl (plus net charge neutralizing ions) for a total of 160,358 atoms. We used Mg^2+^ and GTP parameters as described previously and the CHAMRM36m force field^70^. The initial equilibration simulation was performed for 30 ns with a 2 fs time step using NAMD/2.12 package^69^. Standard parameters were used with Particle-Mesh Ewald (PME) method for long range electrostatic interaction, a cut-off of 1.2 nm for the van der Waals (vdW) potential and short electrostatic interaction, SHAKE algorithm for constraint of all covalent bonds to hydrogen, and a Langevin thermo- and baro-stat at 310 K and 1 bar were used, respectively. The equilibrium simulations were then transferred to the Anton 2 supercomputer for production simulations of 1μs^71^.

### Dynamical Network Analysis

Dynamical network analysis is a general method used to obtain an accurate picture of network topology and long-range signaling in protein complexes derived from molecular dynamics simulations^29^. Each amino acid residue in the complex is assigned a node centered on its Cα atom and used as a base to construct significant regions of amino acid interactions and pathways of allosteric modulation that connect them. Edges are placed to connect the nodes between residues that remain within 4.5 Å distance for at least 75% of the simulation time. The edges are weighted using pairwise correlation data calculated by the program Carma^30^. This information can then be mined to define community networks using the Girvan-Newman algorithm^72^ and the diversity of paths that connect sites of functional significance in the complex. Nodes in the same community network can communicate with each other easily through multiple paths, whereas those in distinct community networks either do not communicate well or communicate through one or a small number of nodes essential for allosteric modulation. Once “source” and “sink” residues are defined as those whose allosteric connections are being evaluated, the shortest allosteric path between them is termed the optimal path. All others are labelled as suboptimal paths. A length offset of 20 edges that prevents the suboptimal paths from being more than 20 edges longer than the optimal path was used to prevent the program from identifying uneccesarily long paths connecting the residues.

## Acknowledgments

Thanks to Andrew Stephen and the Ras initiative team at the Frederick National Laboratory for Cancer Research, NCI, for providing the farnesylated and methylated full-length KRas4B used in the SLB experiments. We thank Richard Gillilan (CHESS) for assistance with collection of the SEC-SAXS data. The protein structure coordinates used in this work were previously published in the PDB and are accompanied in the text by their respective accession codes. This work was supported by NSF grant MCB-1517295 awarded to CM.

