## Supplemental Figures for "Raf promotes dimerization of the Ras G-domain with increased allosteric connections"

### Supplementary Materials

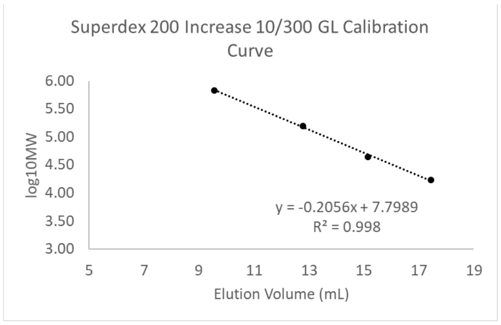


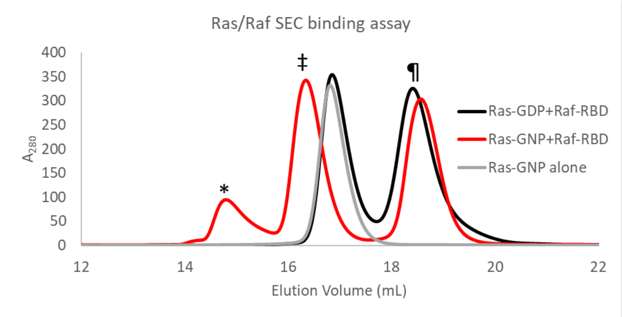


| Protein | Molecular Weight (MW) (Da) | $log_{10}$(MW) | Elution Volume (mL) |
|---|---|---|---|
| Thyroglobulin | 670000 | 5.83 | 9.57 |
| γ-globulin | 158000 | 5.20 | 12.79 |
| Ovalbumin | 44000 | 4.64 | 15.15 |
| Myoglobin | 17000 | 4.23 | 17.44 |

| Protein | Theoretical MW (Da) | $log_{10}$(MW) | Elution Volume (mL) | Calculated MW (Da) |
|---|---|---|---|---|
| Dimer Complex * | 57790.04 | 4.76 | 14.80 | 57127.13 |
| Monomer Complex ‡ | 28895.02 | 4.46 | 16.34 | 27477.98 |
| Raf-RBD ¶ | 9187.74 | 3.96 | 18.57 | 9583.51 |

Fig. S1: Calculation of species molecular weight from SEC elution volume. A Superdex 200 Increase 10/300 GL gel filtration column (GE Healthcare Life Sciences) was used to achieve separation based on molecular weight of the species in solutions containing KRas and CRaf-RBD. A standard curve was constructed using a buffer system and protocol identical to those used in the KRas/Raf-RBD SEC experiment. The elution volumes of the peaks were applied to the calculated standard curve regression to obtain each peak's molecular weight, shown under the column labelled "Calculated MW (kDa)". The theoretical molecular weights were obtained from the known molecular weights of the truncated KRas G-domain and CRaf-RBD and shown in the "Theoretical MW (kDa)" column. The calculated and theoretical molecular weights of each species are in good agreement. The SEC run in red shows formation of the dimer (*) when KRas is bound to the GTP analogue GppNHp (denoted as GNP) and in solution with Raf-RBD. SEC runs containing Ras-GDP and Raf-RBD (black) do not show the dimer peak, as Raf-RBD binds specifically to the GTP-bound form. RasGNP in the absence of Raf-RBD (gray) shows the monomer but not the dimer peak, attesting to the importance of Raf-RBD in dimerization.

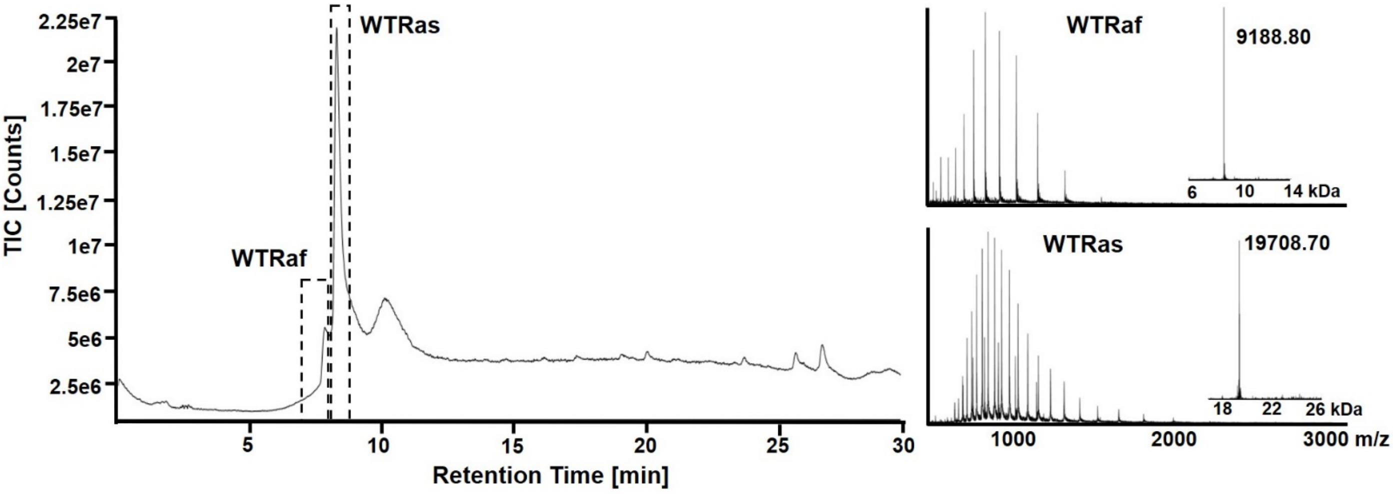


**Fig S2.** Mass spectrometric analysis of the SEC eluent corresponding to the Ras/Raf-RBD complex dimer peak. Intact protein LC-MS was performed using an H class Acquity Ultra High Pressure Liquid Chromatography (UPLC) system coupled with a Xevo G2-S Q-TOF mass spectrometer (Waters Corp, Milford, MA) as described in Donnelly DP 2019 (main text ref 67) Briefly, reversed phase chromatography was employed for separation (Acquity UPLC protein BEH C4 300 Å pore size, 1.7 µm particle size, 100 mm bed length, 2.1 mm ID x 100 mm) with 95% water/ 5% acetonitrile with 0.1% formic acid as solvent A and 95% acetonitrile/ 5 % water with 0.1% formic acid as solvent B. The reversed phase chromatogram is shown on the left and the mass spectra on the right. Within each mass spectrum is an inset showing the maximum entropy deconvolution of the time average spectrum, which are consistent with the theoretical molecular masses of Raf (top) and Ras (bottom). The third peak has the same mass as WT Ras-RBD, ruling out this peak arising from a disulfide bridged, or any other covalent linkage, multimer.

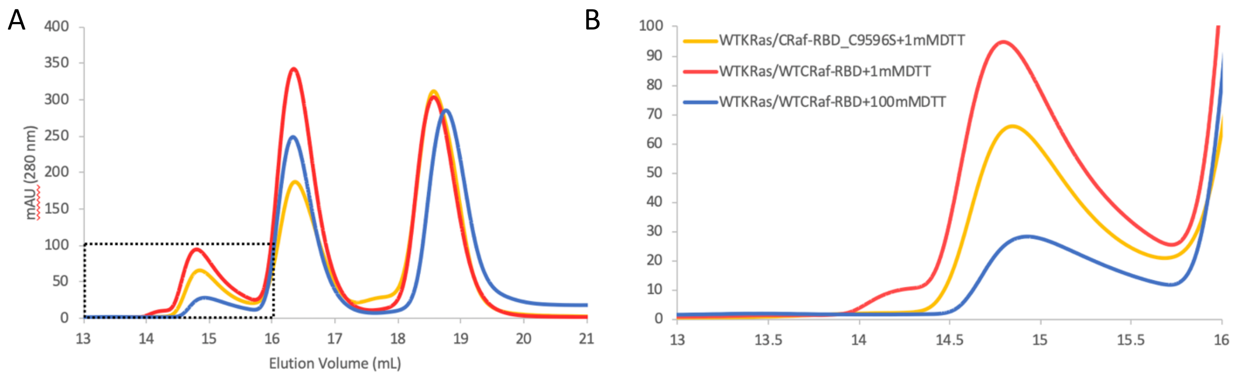


Fig. S3: SEC runs for KRas and Raf-RBD showing the presence of dimers without disulfide bonds. (A) Full run showing the expected peaks for the dimer of the KRas/Raf-RBD complex, the monomer of the KRas/Raf-RBD complex and Raf-RBD on a Superdex 200 Increase 10/300 GL gel filtration column (GE Healthcare Life Sciences). (B) Enlargement of the dimer peak, shown in the boxed section in A. The red curve is the same as shown in Fig. S1, which represents KRas and wild-type Raf-RBD in the presence of 1mM DTT. The yellow cure is for KRas and Raf double mutant C95S-C96S with 1mM DTT. The blue curve is for KRas and Raf-RBD in 100 mM DTT. There is variation in the relative amount of dimer present, as mentioned in the main text. Note the small peak of higher MW that appears before the dimer peak for the run using wild type Raf-RBD with 1 mM DTT (red curve). This peak is not present with either the Raf-RBD C95S-C96S double mutant in 1mM DTT or when 100 mM DTT is used with the wild type Raf-RBD, indicating that the high MW shoulder is likely due to disulfide bonds between exposed cysteine residues in the dimer forming larger order complexes.

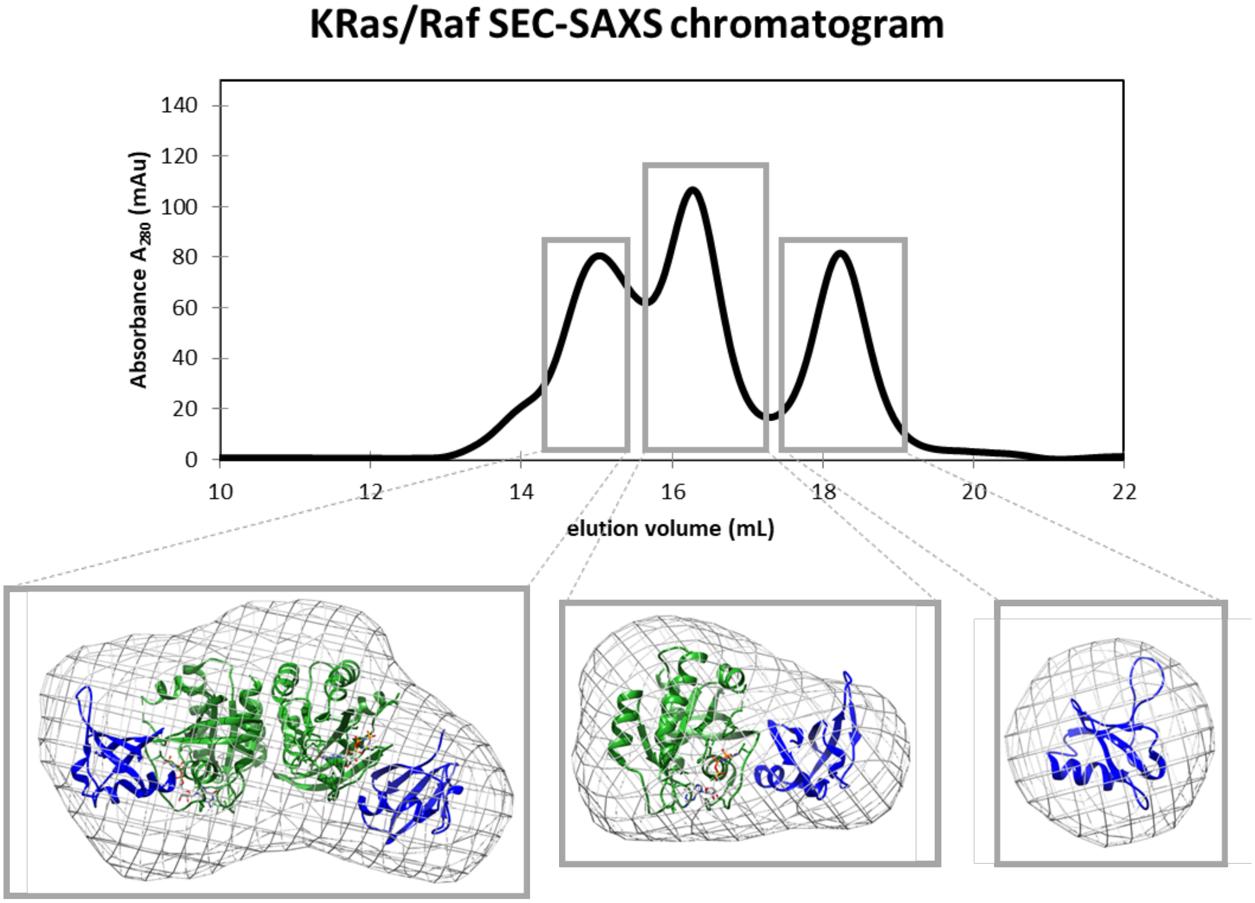


Fig. S4: Calculated envelopes of the three dominant species resulting from the SEC-SAXS experiment for KRas-GppNHp (truncated at residue 166) in the presence of excess Raf-RBD. The three major peaks eluting from a Superdex 10/300 Gel Filtration column (GE Healthcare Life Sciences) were subjected to in-line SAXS data collection and envelope reconstruction using BioXTAS RAW (*35*). SAXS data obtained from the peak corresponding to the dimer species has a calculated radius of gyration ($R_g$) of 30 Å. The dimer of the complex obtained from our crystal structure of the HRas/CRaf-RBD complex (PDB ID 4G0N with a two-fold symmetry rotation axis applied to generate the dimer) fits the SAXS data with a $\chi^2$ value of 1.24. The second species has an $R_g$ of 21 Å, and the monomer of the HRas/CRaf-RBD complex (PDB ID: 4G0N) fits the data with a $\chi^2$ value of 1.01. Raf-RBD (PDB ID: 1RRB) alone has the smallest $R_g$, 13.3 Å, and calculated fit to the SAXS data with a $\chi^2$ value of 0.92. The obtained $\chi^2$ values are all within the range expected for excellent fit of the models to the data. Chi squared values of the theoretical model and experimental scattering correlations were calculated using the FOXS server (main text ref 66).

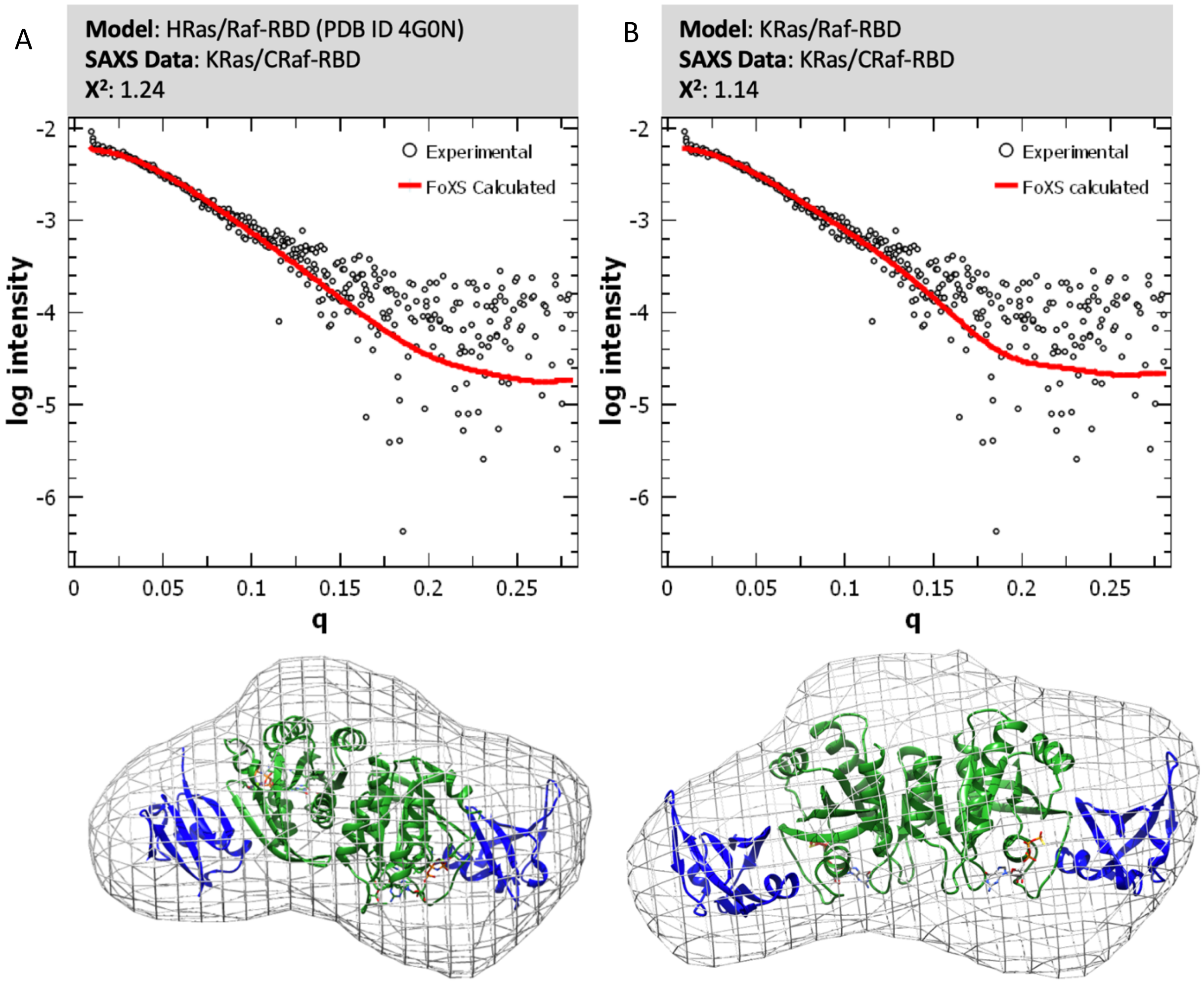


Fig. S5: Quantitative analysis of crystallographic and NMR-based dimer models against SAXS data using FOXS (main text ref 66). The SAXS data used was that obtained for the dimer peak eluted from a SEC column (KRas was truncated at residue 166). FOXS generates a theoretical scattering curve from a given model (red on upper portion of the panels) and provides a match to the experimental SAXS data. The fit of the calculated scattering curve for the rigid model to the experimental X-ray scattering data is given by the $\chi^2$ value, with values less then 2.0 indicative of a good fit. (A) The HRas/CRaf-RBD dimer model was generated from symmetry related molecules in the 4G0N crystal structure. (B) The KRas/Raf-RBD dimer model was obtained from an average of the top 20 NMR models published by Lee et al (11). The average was calculated using VMD (main text ref 68). As this model does not include Raf-RBD, the Raf-RBD domain was placed in each monomer in this model via superposition on the HRas/Raf-RBD complex (PDB ID: 4G0N). The resulting $\chi^2$ values indicate excellent fits to the SAXS data for both models ($\chi^2$ values < 1.5).

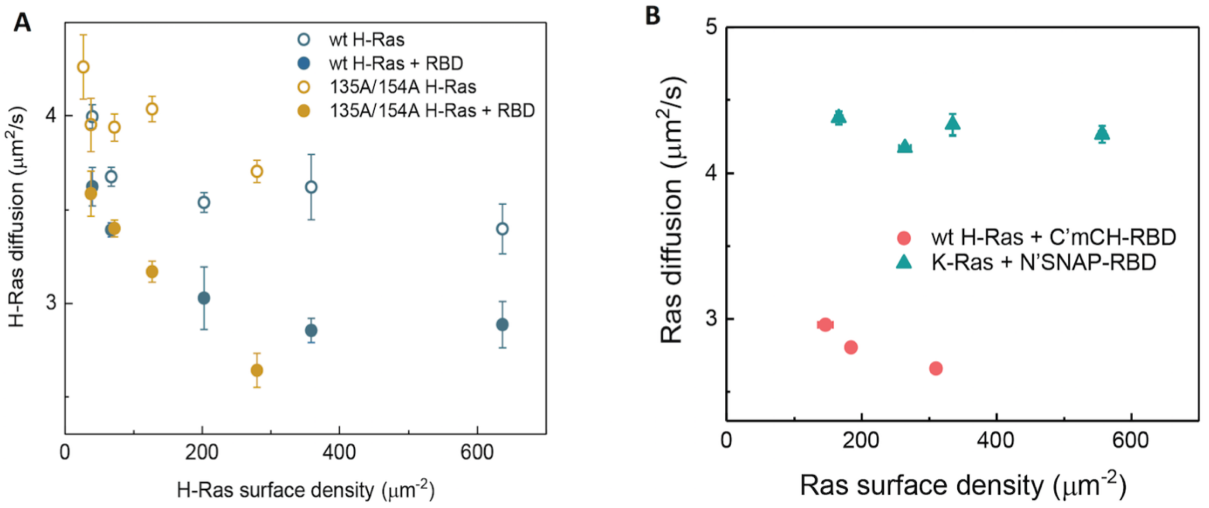


Fig. S6. (A) FCS diffusion measurements for wild-type HRas and dimer interface mutant at the α4-α5 interface (R135A/D154A) with and without 250 nM Raf-RBD, showing identical complexation behavior. Interestingly, the double interface mutant HRas R135A/D154A has diffusion behavior identical to the wild type. This could be an indication that mutation of the two residues to the small alanine side chain is tolerated in the presence of the HVR and the membrane, given an otherwise intact Ras dimer interface. This behavior is contrary to that observed for the D154Q substitution shown to completely disrupt KRas dimerization *in vitro* and in cells (main text ref 16) and for the R135E charge reversal substitution that disrupts Ras/Ras interactions measured by paramagnetic relaxation enhancement NMR (main text ref 12), possibly due to steric hindrance from the bulky mutant side chains at the interface. (B) Modification of Raf-RBD results in different oligomerization behavior. mCherry at the C-terminus preserves the oligomerization, while SNAP tag at the N-terminus compromises it.

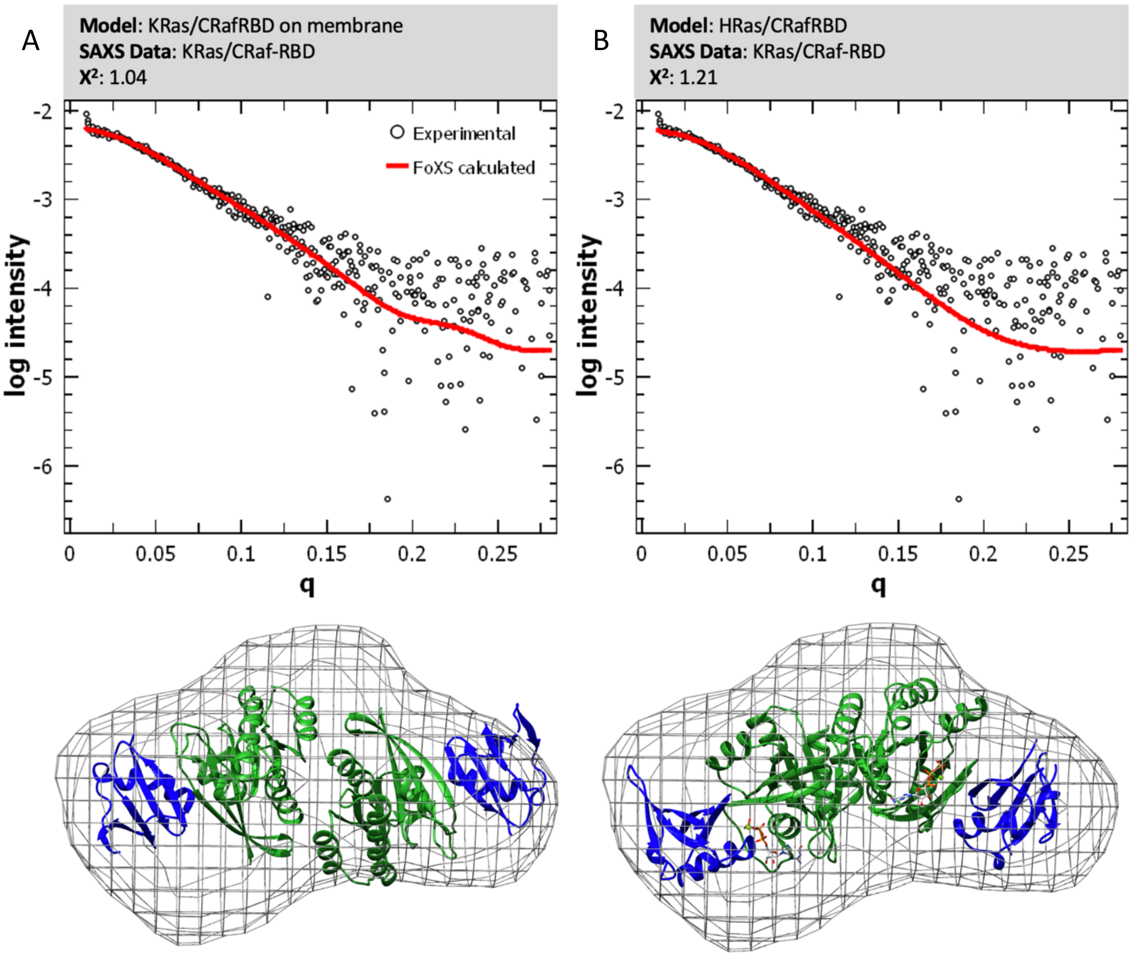


Fig. S7: Quantitative analysis of KRas/CRaf-RBD and HRas/CRaf-RBD dimer models against SAXS data using FOXS (main text ref 66). The same SAXS data was used as for Fig. S5. (A) The KRas/CRaf-RBD dimer model was generated from the average structure resulting from 1 μs simulation of the full-length lipidated KRas in complex with CRaf-RBD on the membrane, with KRas truncated at residue 166 to match the Ras construct used for SAXS data collection. (B) The HRas/CRaf-RBD dimer model was generated from the average structure of the 90 ns simulation of the crystallographic dimer (PDB ID 4G0N). The resulting $\chi^2$ values for both dimer models to the SAXS data indicate excellent fit for both models ($\chi^2$ values < 1.5).

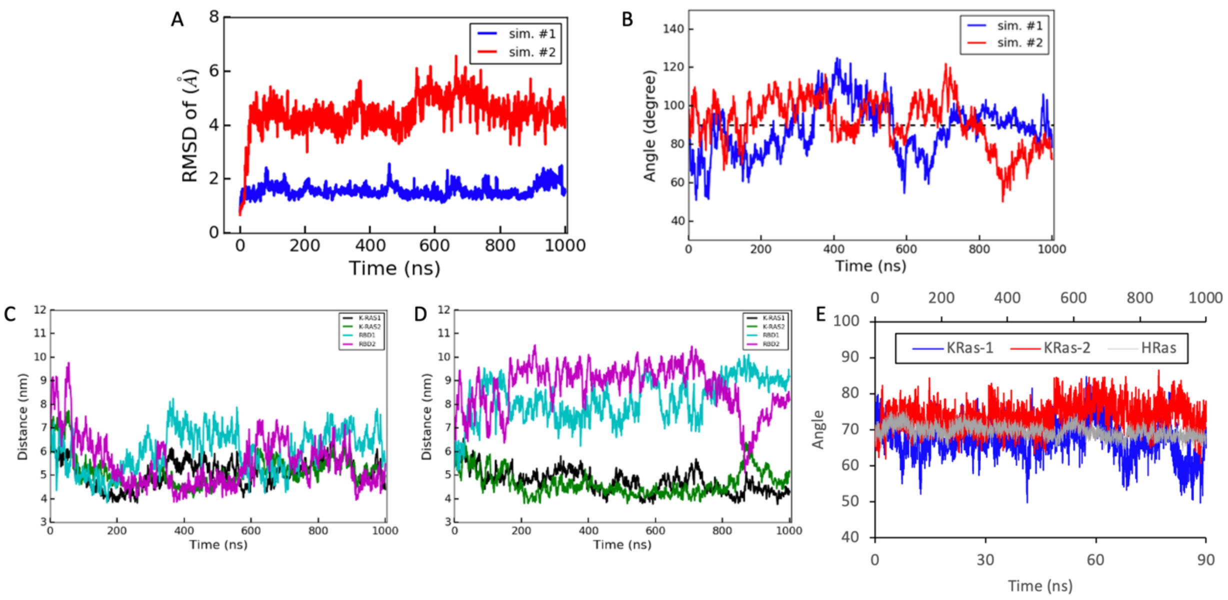


Figure S8. Properties of the 1 μs simulations of the KRas/Raf-RBD dimer on the membrane. (A) Root mean square deviation (RMSD) from the starting model of the KRas/Raf-RBD dimer throughout the two simulations. (B) Angle of the KRas/KRas dimer relative to the membrane in the presence of Raf-RBD. The angle is measured between a vector normal to the membrane and a vector connecting the centers of the two KRas catalytic domains. A dashed line indicates 90°, with the Ras helices perpendicular to the plane of the membrane. (C) and (D) show the distance from the center of mass of each component of the KRas/Raf-RBD dimer to the membrane center over the course of the first and second simulations, respectively. (E) Angle between helices 4 and 5 across the dimer interface for 90 ns of the two simulations with KRas (KRas-1 blue, KRas-2 red) compared to the 90 ns simulation with HRas (gray). The angle was defined as that between the vectors connecting the center of mass calculated for residues at the two ends of the dimer interface helices in each Ras protomer. At the end furthest from the membrane, the center of mass was calculated from helix 4 residues 127, 128, 129, 130 and helix 5 residues 152, 153, 154, 155. At the end closest to the membrane helix 4 residues 134, 135, 136, 137 and helix 5 residues 163, 164, 165, 166 were used. The average angle sampled by each isoform differs by less than 10°. However, the KRas angle fluctuates in the range of 20-30° around the average, while the angle in HRas stays within 10° of the average.
